# Mechanically Matched Silicone Brain Implants Reduce Brain Foreign Body Response

**DOI:** 10.1101/2020.12.13.422566

**Authors:** Edward N. Zhang, Jean-Pierre Clément, Alia Alameri, Andy Ng, Timothy E. Kennedy, David Juncker

## Abstract

Brain implants are increasingly used to treat neurological disorders and diseases. However, the brain foreign body response (FBR) elicited by implants affects neuro-electrical transduction and long-term reliability limiting their clinical adoption. The mismatch in Young’s modulus between silicon implants (∼180 GPa) and brain tissue (∼1-30 kPa) exacerbates the FBR resulting in the development of flexible implants from polymers such as polyimide (∼1.5-2.5 GPa). However, a stiffness mismatch of at least two orders of magnitude remains. Here, we introduce (i) the first mechanically matched brain implant (MMBI) made from silicone (∼20 kPa), (ii) new microfabrication methods, and (iii) a novel dissolvable sugar shuttle to reliably implant MMBIs. MMBIs were fabricated via vacuum-assisted molding using sacrificial sugar molds and were then encased in sugar shuttles that dissolved within 2 min after insertion into rat brains. Sections of rat neocortex implanted with MMBIs, PDMS implants, and silicon implants were analyzed by immunohistochemistry 3 and 9-weeks post-implantation. MMBIs resulted in significantly higher neuronal density and lower FBR within 50 µm of the tissue-implant interface compared to PDMS and silicon implants suggesting that materials mechanically matched to brain further minimize the FBR and could contribute to better implant functionality and long-term reliability.

## 1. Introduction

Brain implants have significant implications for the treatment of neurological disorders and diseases. Challenges, however, in long-term reliability and maintenance of high-quality recordings have prevented the widespread clinical adoption of brain implants.^[1]^ The brain foreign body response (FBR) elicited by implants affects their functionality due to the formation of dense glial scars that isolate the implants from neurons of interest and persistent inflammation leading to neuronal death around the implants.^[2]^ The mismatch in Young’s modulus (*E)* between current implants made from rigid materials such as silicon (∼180 GPa)^[3]^ and brain tissue (∼1-30 kPa)^[4-6]^ exacerbates the FBR due to strain at the tissue-implant interface^[7]^ and the response of glial and neuronal cells to stiff materials.^[8, 9]^ Consequently, significant efforts have been devoted to developing softer implants made from polymers such as Parylene-C (∼3-4 GPa),^[6,10-12]^ SU-8 (∼2 GPa),^[12]^ polyimide (∼1.5-2.5 GPa),^[12-15]^, and PDMS (∼1-1.6 MPa).^[15, 16]^ However, current compliant implants are still two or more orders of magnitude stiffer than the brain.

Implants with hydrogel coatings that mechanically match brain tissue have been developed recently,^[17]^ and while they are mechanically matched at the tissue-implant interface, they cannot mask strain owing to larger implant movements. An alternative strategy has been to make thinner implants, some at sub-micron scale.^[18]^ This imparts flexibility to even rigid materials resulting in reduced bending stress and FBR. However, this approach is not effective in reducing axial and stretching strain that could arise under conditions such as brain swelling or significant rotational displacement of the brain. Conceptually, mechanically matched brain implants (MMBIs) would reduce FBR significantly. However, whether this is true is unknown as a MMBI has yet to be made. Ascertaining whether MMBIs induce appreciably lower FBRs compared to current complaint implants will help answer if efforts in developing them is worthwhile.

The realization of MMBIs faces fabrication and implantation challenges. Soft micro-elastomeric devices are commonly made by replica molding, but ultra-soft materials are difficult to release from molds because they are tacky and often get damaged during release. Consequently, sacrificial molds have been introduced for the fabrication of ultra-soft devices.^[19, 20]^ Replica molding also suffers from residual membranes forming on the surface of the molds. These membranes caused by the overfilling of prepolymer prevent the formation of independent device features. Current solutions such as cure inhibition^[21]^ or physical expulsion of the excess prepolymer^[22]^ and etching of the membrane^[23]^ results in rough or sticky surfaces.

MMBIs require a temporary increase in structural strength in order to penetrate brain tissue for implantation. Whereas numerous strategies to aid the insertion of flexible implants exist, they generally entail the use of dissolvable coatings or temporary rigid shuttles. However, many of these strategies are incompatible with MMBIs due their soft and tacky nature. For example, affixing MMBIs to rigid shuttles using dissolvable adhesives^[6, 16]^ or electrostatic interactions^[15]^ can result in unreliable decoupling of MMBIs from the shuttles due to residual adhesion and hydrophobic interactions. Unsuccessful decoupling can cause inaccurate implant positioning in the brain or the implant being pulled out from the brain during shuttle removal.^[15]^

Dissolvable coatings circumvent the need for shuttle removal. Materials such as sugar,^[16]^ polyethylene glycol (PEG),^[6]^ tyrosine-derived polymers,^[12]^ and carboxy-methyl-cellulose (CMC)^[11]^ have been used. However, most of these materials suffer from significant drawbacks. Tyrosine-derived polymers have relatively long dissolution times (> 1 h),^[12]^ while CMC becomes a soft gel *in vivo* rather than dissolving.^[11]^ PEG, which is often used to stiffen flexible implants, has limited rigidity and fails to provide the structural strength for MMBIs.^[12]^ Furthermore, these materials are often applied by dip-coating making the thickness and shape difficult to control.^[13, 14]^ Micro-molding offers better control of the shape but is susceptible to overflow of the coating material that requires manual removal leading to variations in coating thicknesses.^[11]^ Micromoulding in capillaries (MIMIC)^[12]^ avoids overflow but is limited to low viscosity solutions (< 400 mPa.s),^[24]^ which are not compatible with more viscous materials like sugar-based solutions.

Here, we introduce the first MMBIs fabricated using vacuum-assisted molding (VAM) in sacrificial sugar molds. VAM is also used to encase the MMBIs in sugar shuttles that dissolve quickly upon insertion into rat brains. To study the effects of implant stiffness on the FBR, PDMS (∼1.6 MPa) and silicon (∼180 GPa) implants of the same dimensions were also fabricated and inserted into rat brains using dissolvable sugar shuttles for comparison. Immunohistochemical analyses of cortical sections from adult rat brains sacrificed at 3 and 9-week post-implantation were conducted to assess the FBR and neuronal density surrounding the tissue-implant interface of the different implants.

## 2. Results and Discussion

### 2.1. Mechanically Matched Material for Brain Implant Fabrication

Ecoflex, a platinum-catalyzed silicone elastomer was selected to make MMBIs because of its low stiffness, biocompatibility, and high electrical breakdown strength.^[25, 26]^ The Young’s modulus of Ecoflex was determined from stress-strain curves acquired from tensile tests performed on Ecoflex test specimens fabricated in accordance to the American Society for Testing of Materials D412-C specifications (Figure S5-S6, SI). Since MMBIs were sterilized at 121 °C for 45 min and then encased in sugar shuttles using a vacuum oven set to 240 °C for 20 min, we investigated how these high temperature processes affected the stiffness of MMBIs. We found no statistical difference between the stiffness of Ecoflex before and after being autoclaved and heated in a vacuum oven (∼20 kPa, p-value = .67). For comparison, the stiffness of PDMS before and after undergoing the high temperature processes increased significantly from 1.07 ± .02 to 1.60 ± .05 MPa (p-value < .001). This indicates that Ecoflex is thermally stable after curing at 60 °C for 1 h and demonstrates superior thermal stability compared to PDMS cured at 60 °C for 24 h. The stiffness of Ecoflex MMBIs is compared to other compliant implants in **Figure 1**.

**Figure 1.**
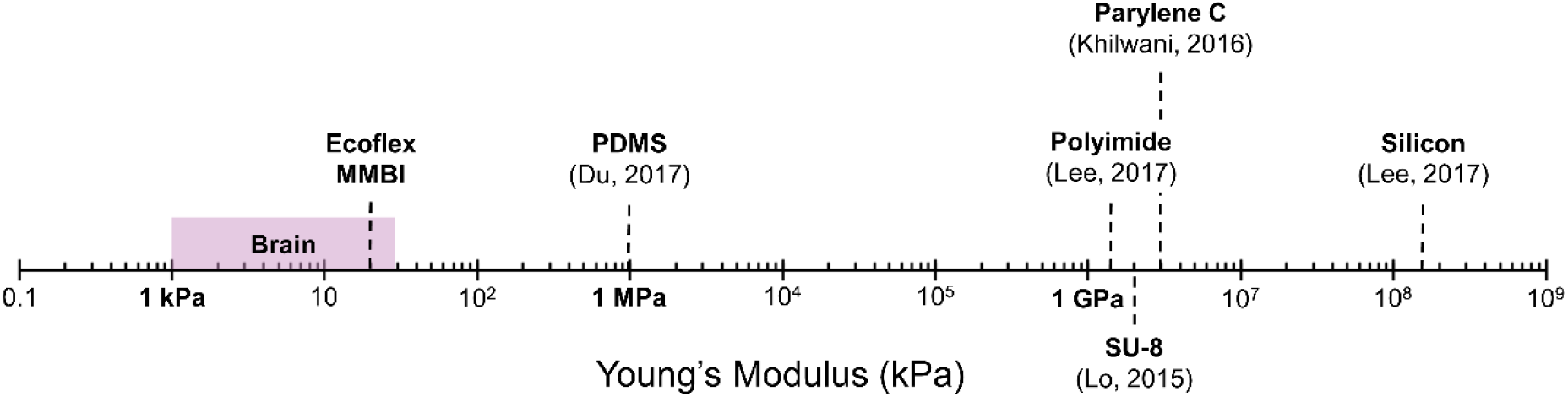
Logarithmic graph showing the Young’s modulus of the brain and various materials used for making brain implants. MMBIs are the softest implants developed to date to the best of our knowledge.

### 2.2. Fabrication of Sacrificial Sugar Molds

The realization and implantation of MMBIs required three distinct processes: fabrication of sacrificial sugar molds, VAM of MMBIs in sugar molds, and encasing MMBIs in micro-molded dissolvable sugar shuttles. Using sugar molds that dissolved in water circumvented the need to release MMBIs manually, preventing potential damage and distortion. **Figure 2a** illustrates the sugar mold fabrication process. Positive SU-8 master molds with MMBI features that were 300 µm wide, 200 µm thick, and 5 mm long (Figure 2b) were fabricated using conventional photolithography. The dimensions chosen are commensurate with existing implants and were chosen to facilitate analysis of implant stiffness on the FBR. However, the dimensions could be reduced towards further minimizing tissue damage and local perturbation by creating a master mold with smaller features.

**Figure 2.**
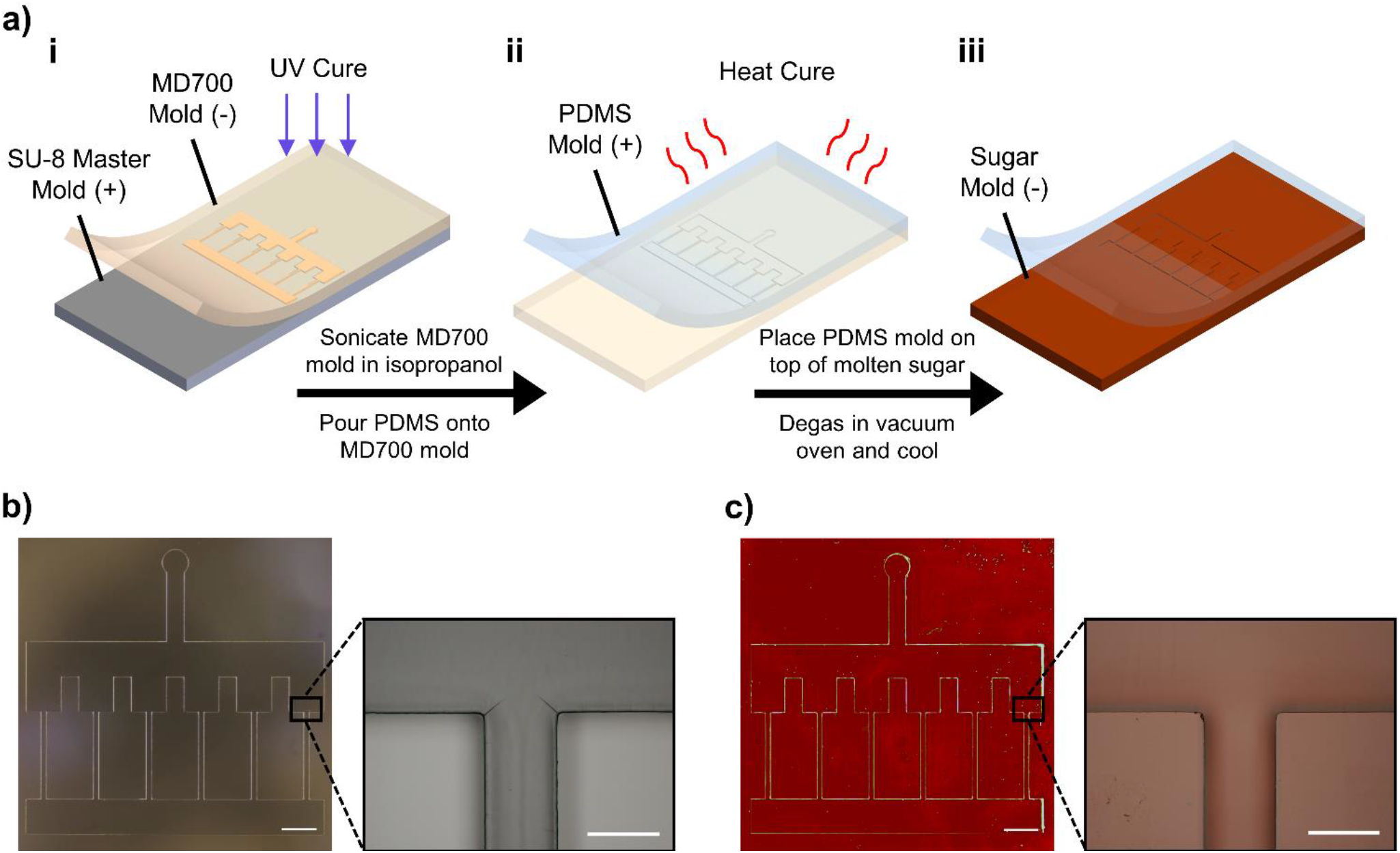
Fabrication process of sacrificial sugar mold. a-i) MD700 prepolymer is UV cured on top of positive SU-8 master mold to create negative MD700 mold. a-ii) PDMS prepolymer is heat cured on top of MD700 mold to create positive PDMS mold. a-iii) PDMS mold is released from solidified sugar mold. b) Microscope image of positive SU-8 master mold. c) Microscope image of negative sugar mold. Scale bars: 2 mm, 300 μm (insets).

MD700, a bifunctional perfluoropolyether-urethane methacrylate, prepolymer was UV cured on top of the SU-8 molds to create negative molds. MD700’s low surface energy (∼15 mN/m)^[27]^ allowed for easy release from SU-8 molds without needing surface treatments such as silanization, circumventing potential contamination of the sugar molds with cytotoxic chemicals. Positive PDMS molds replicated from the MD700 molds were used to create the sugar molds. PDMS’s high thermal stability^[28]^ allowed the mold to withstand the high processing temperatures of the sugar molds and its low surface energy (∼30 mN/m)^[29]^ prevented it from sticking to the sugar molds.

A mixture of light corn syrup, water, and sucrose in a 1:2:3 ratio by volume was used to make the sugar molds. Sugar mixtures typically reach the hard crack stage, the point where once cooled become hard, at ∼150 °C.^[30]^ However, sugar mixtures heated to 150 °C have a relatively low glass transition temperature (*T*_g_) of ∼30 °C.^[19]^ Since the Ecoflex prepolymer used to fabricate MMBIs was cured at 60 °C, sugar mixtures were heated to 180 °C to raise the *T*_g_ of the resultant sugar molds. Extending the caramelization process polymerizes more monosaccharide molecules in the sugar mixtures raising the *T*_g_ of the sugar molds beyond 60 °C.^[31]^ PDMS molds were placed feature-side into a silicone beaker containing the molten sugar and then degassed in a vacuum oven set to 240 °C (∼150 °C sugar temperature) to remove bubbles from the molten sugar that would compromise the features of the sugar mold. After degassing, the molten sugar was cooled at room temperature (RT), and once solidified, the PDMS molds were released to create the sugar molds. A comparison of a SU-8 mold and a sugar mold (Figure 2c) shows high-fidelity replication of the SU-8 mold features.

We found sugar molds left at RT with a relative humidity of ∼40% begin to degrade within 1 h due to sugar’s hydroscopic nature. Storing them in a vacuum desiccator, however, kept the molds preserved for over a month. Sugar is highly advantageous for fabricating sacrificial molds due to its relatively high stiffness of ∼100 MPa,^[19]^ which allows for high-fidelity replication of features. Additionally, sugar is inexpensive, highly soluble in water, and non-toxic.

### 2.3. Fabrication of Ultra-Soft Brain Implants

Here we adapt VAM, which was previously developed in our lab for making polymer microfilter membranes,^[32]^ to mold MMBIs using Ecoflex. Pouring prepolymer into the molds leads to residual membranes due to overflow (Figure S3, SI), which is circumvented by VAM. **Figure 3a** illustrates the sugar mold setup for VAM and the fabrication process of MMBIs. Heat release tape (HRT) was used to seal the sugar mold while leaving the inlet accessible, and a PDMS O-ring was placed around the inlet to contain the Ecoflex prepolymer. When degassed in a vacuum desiccator, air inside the mold escaped by forming air bubbles in the prepolymer (Figure 3b). The desiccator was vented once the bubbles subsided (∼20 min), and the difference between the ambient pressure and the pressure inside the mold caused the Ecoflex prepolymer to infill the mold within seconds. VAM of Ecoflex is time sensitive because Ecoflex has a pot life of 30 min whereby its viscosity doubles. We observed that sometimes infilling failed because the prepolymer viscosity became too high after degassing. To lower the prepolymer viscosity, a biocompatible silicone fluid was added to the mixture. The prepolymer was kept on ice after mixing until use for VAM. After curing, the MMBIs were released within minutes by dissolving the molds in water. MMBIs accurately replicated the mold features and were trimmed in half using a razor blade to form 2.5 mm long MMBIs.

**Figure 3.**
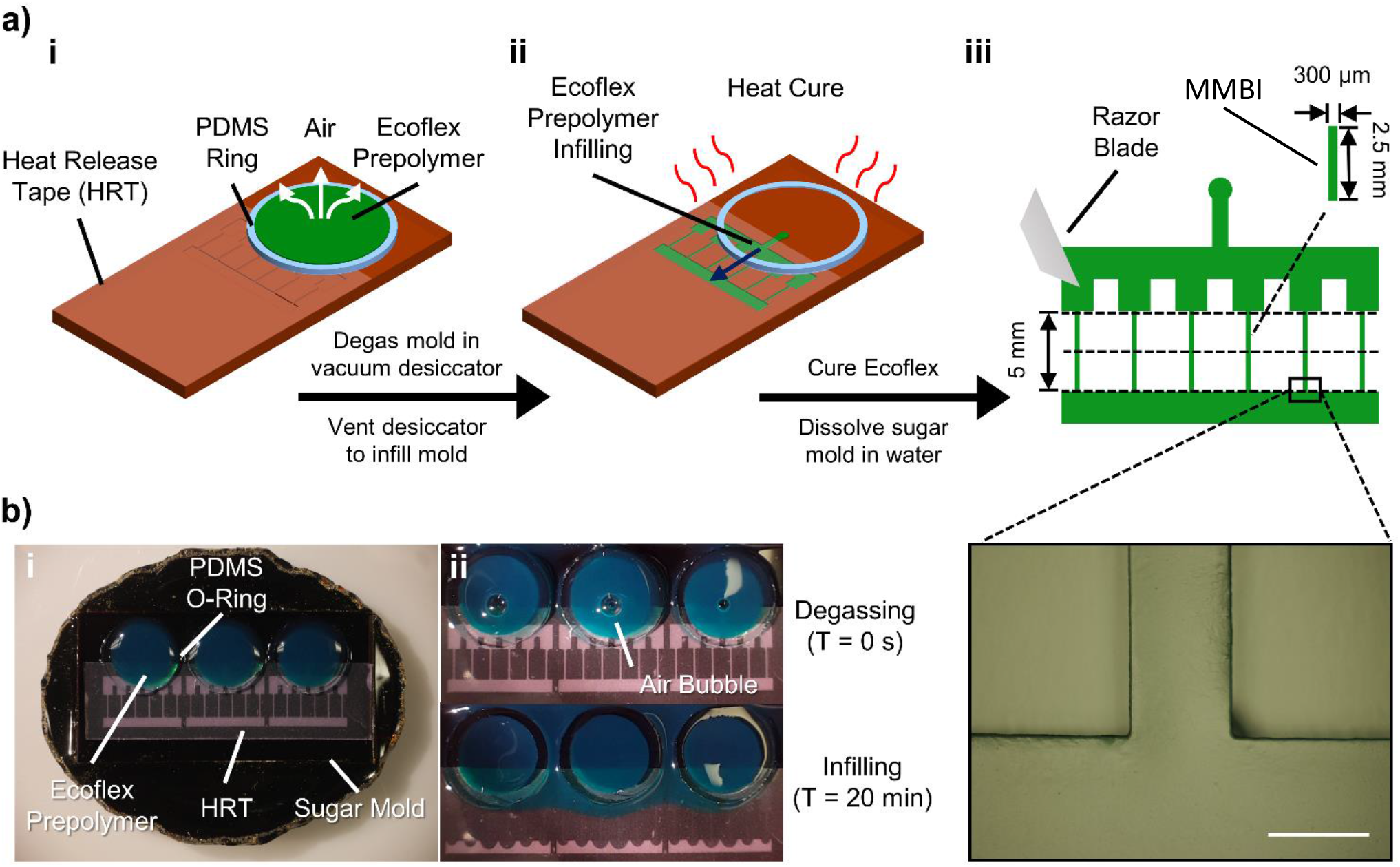
Fabrication process for MMBIs. a-i) Sugar mold is sealed with heat release tape (HRT) and Ecoflex prepolymer is pipetted into PDMS O-ring. Mold is degassed. a-ii) Desiccator is vented and prepolymer infills mold. a-iii) HRT is removed after Ecoflex cures and sugar mold is dissolved in water releasing MMBIs. A razor blade is used to trim 2.5 mm long MMBIs. Microscope image of a molded USBI shown in inset. b-i) Sugar mold setup for vacuum-assisted molding. b-ii) Time-lapse of VAM. Air escapes from mold through prepolymer during degassing indicated by air bubbles. Infilling happens within seconds after desiccator is vented. Scale bar: 300 μm.

VAM’s infilling mechanism whereby vacuum draws in the prepolymer into the mold without active injection could be particularly advantageous for branched structures such as high-density electrode arrays, which could be incorporated into a functional MMBI. Prior to developing VAM, we observed delamination of mold seals when injecting Ecoflex into molds with features in the hundreds of micrometers using a syringe pump due to the high injection pressure. For molds with arrays of features at the scale of tens of micrometers, active injection becomes an even greater challenge. VAM coupled with sacrificial sugar molds is an effective and reliable method of fabricating soft micro-elastomeric devices with complex designs down to micrometer dimensions.

### 2.4. Encasing MMBIs in Dissolvable Sugar Shuttles

MMBIs cannot be directly implanted into the brain. Hence, we encased MMBIs into dissolvable sugar shuttles fabricated using VAM (**Figure 4a)**. The shuttle design faces a trade-off between larger dimensions to provide structural integrity and smaller dimensions to minimize tissue damage. Here, shuttles were 700 μm wide, 450 μm thick, and 8 mm long. Master molds with these shuttle dimensions were 3D-printed and replicated into negative PDMS molds that were used to fabricate the shuttles. MMBIs, as well as PDMS and silicon implants, were manually inserted into the PDMS molds that were subsequently sealed with a softer PDMS slab with a punched-out hole to expose the mold inlet (Figure 4b). The softer slabs provided superior conformal contact resulting in a more reliable seal. We observed that the MMBIs adhered to the PDMS molds preventing sugar from encasing the side of the MMBIs touching the molds during VAM. Consequently, the released shuttles had hollow cavities without the MMBIs encased. This was resolved by coating the mold features with a thin layer of powdered sugar before inserting the MMBIs into the molds for VAM.

**Figure 4.**
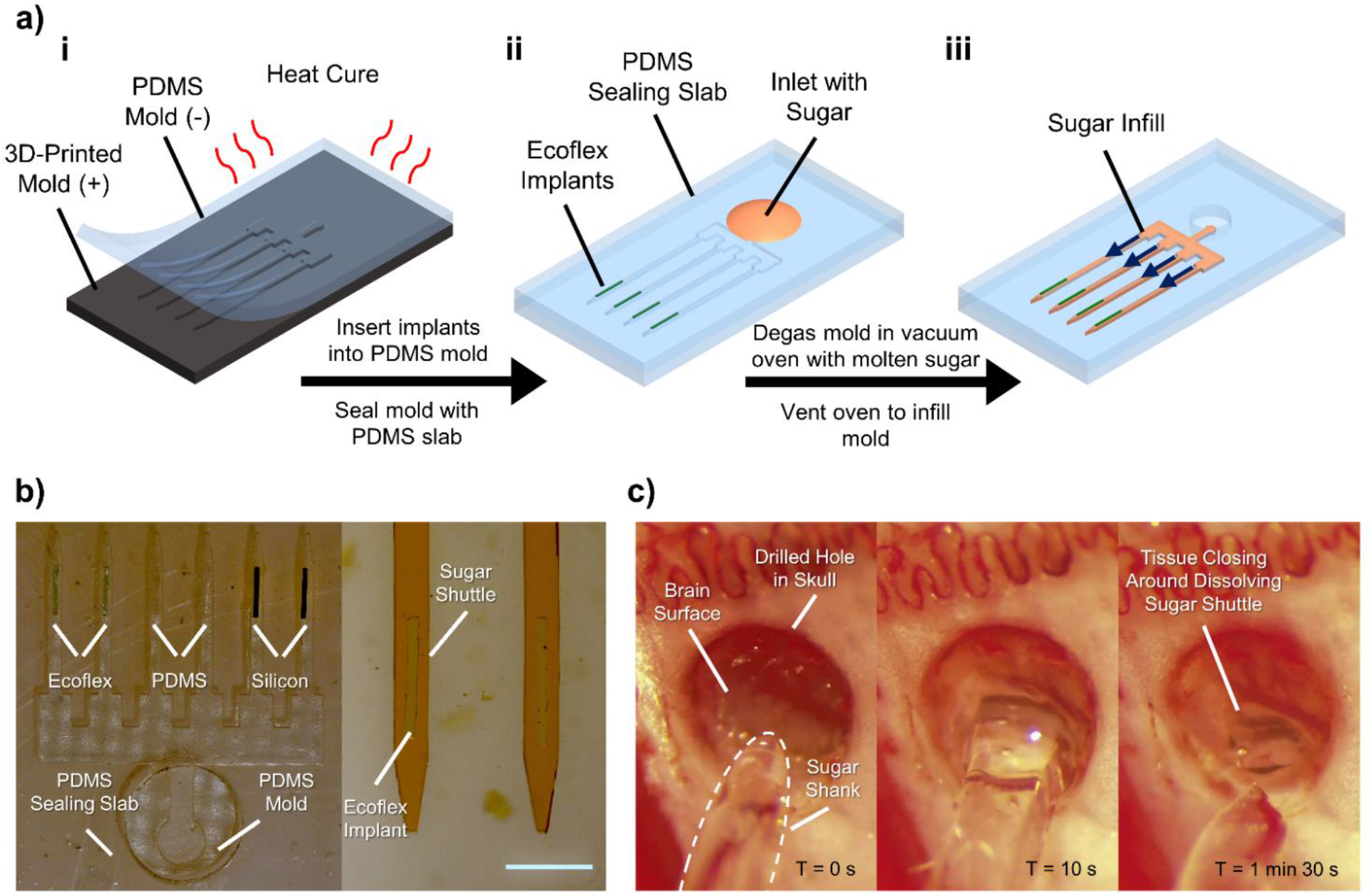
Encasing process for implants in dissolvable sugar shuttles. a-i) Replication of PDMS negative mold from 3D-printed positive master mold. a-ii) Insertion of Ecoflex MMBIs into PDMS mold and sealing mold with PDMS slab for VAM. a-iii) Infilling of molten sugar to encase implants using VAM. b) VAM setup with MMBI, PDMS, and silicon implants embedded into PDMS mold for *in vivo* studies (left). Microscope image of sugar shuttles with MMBIs encased (right). c) Insertion of sugar shuttle with MMBI encased into adult rat brain showing the dissolution of shuttle within 2 min. Scale bar: 1 mm.

The sugar mixture for dissolvable sugar shuttles is identical to the mixture for sugar molds. The sealed PDMS molds were placed with the inlet side downwards into a silicone beaker containing molten sugar heated to ∼150 °C. VAM was done inside a vacuum oven set to 240 °C, which kept the molten sugar at ∼150 °C, to degas the molds and to maintain the molten state of the sugar required for infilling. Following VAM (∼20 min), the infilled sugar was left to solidify. Afterwards, the PDMS sealing slabs were removed, and the resultant sugar shuttles with the encased implants were released from the molds using a tweezer. Release was greatly facilitated by using thin PDMS molds (< 3 mm) that were flexible, which minimized the amount of force required to release the sugar shuttles and achieved a yield of ∼80%. Sugar shuttles with MMBIs encased are shown in Figure 4b. As with the sugar molds, the dissolvable sugar shuttles can be preserved in a vacuum desiccator until use.

The temperature needed to melt the sugar mixture requires that the materials used to fabricate MMBIs must withstand 150 °C. Although this is compatible with metals and many polymers, the high temperature nevertheless restricts the range of materials that can be used to fabricate MMBIs. However, alternative biocompatible materials that do not require high processing temperatures such as silk can be used with VAM to create dissolvable shuttles effectively circumventing this issue. This would enable the incorporation of anti-inflammatory molecules that could be delivered simultaneously to reduce the acute brain FBR and consequently further reducing the chronic FBR. It is important to note that the main advantage of using sugar to create the dissolvable shuttles is that the brain readily metabolizes sugar and it can pass through the blood-brain barrier (BBB).^[33]^ While other biocompatible materials can be used to create the dissolvable shuttles, the long-term effects of materials that are not normally found in the brain or that cannot pass through the BBB are unknown.

### 2.5 Delivery of Dissolvable Sugar Shuttles and Implants into Rat Brains

Figure 4c shows the insertion of a sugar shuttle with an encased MMBI into the neocortex of an adult Sprague Dawley rat. The shuttle dissolves rapidly in less than 2 min leaving the MMBI securely in place, and the tissue displaced by the shuttle during insertion can be seen refilling the implantation site (Video S1, SI). Implants were inserted to reach a target depth of ∼3 mm, slightly deeper than the thickness of the cortex in the rat brain. Sugar shuttles were designed to have a tip angle of ∼17° to minimalize tissue dimpling.^[34]^ The shuttle tip was ∼200 μm in width, corresponding to the resolution limit of the 3D-printer used. Dimpling of the tissue during insertion was effectively minimized by this design. **Figure 5** shows a magnetic resonance image (MRI) of a rat brain sacrificed immediately after surgery depicting the MMBI as a black rectangular shape matching the dimensions of the implanted MMBI confirming accurate positioning, the dissolution of the sugar shuttles, and the refilling of the displaced tissue.

**Figure 5.**
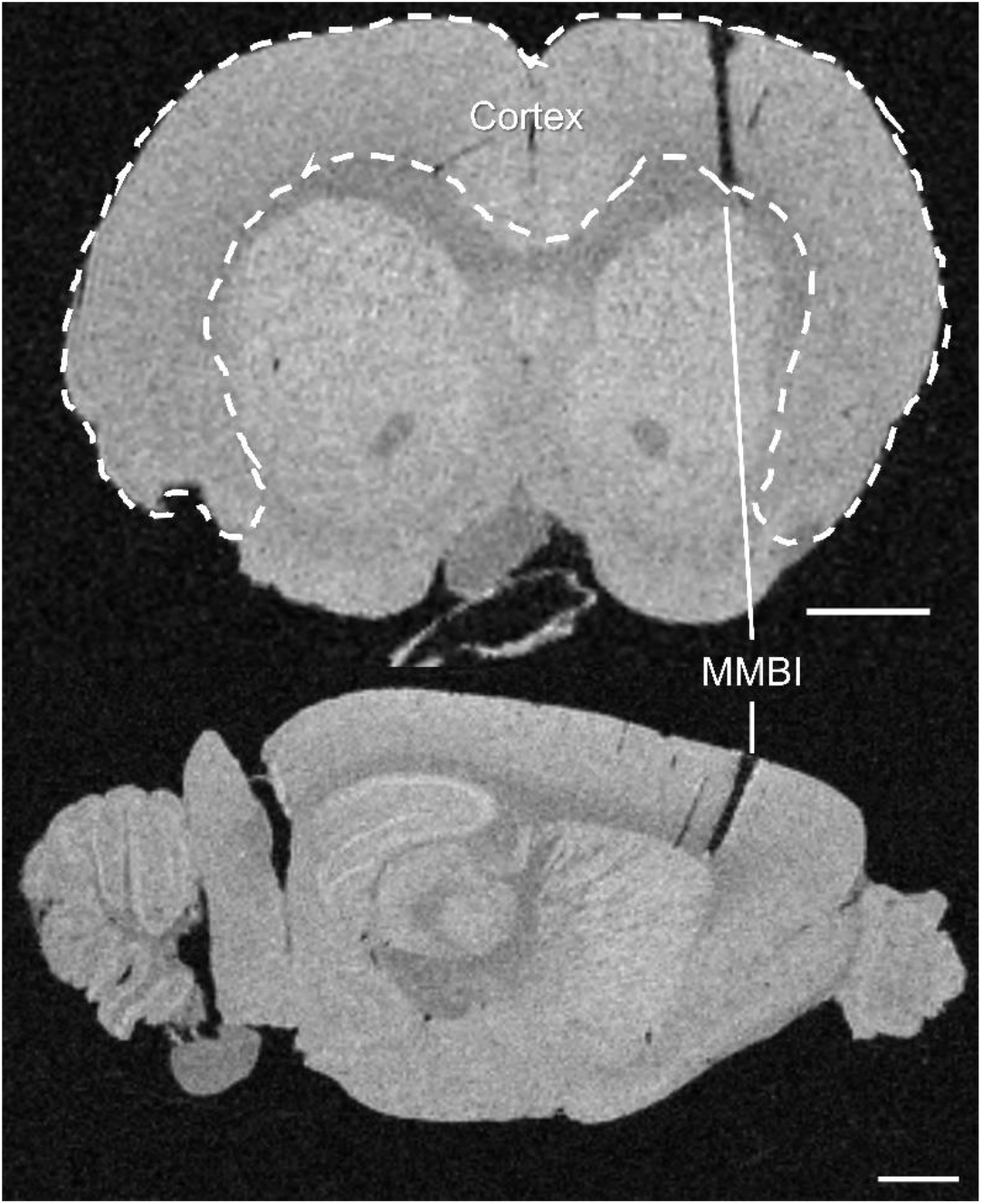
MRI of an adult rat brain sacrificed immediately after surgery showing successful implantation of an MMBI into the cortex delivered using a dissolvable sugar shuttle. Frontal view (top) and sagittal view (bottom). Scale bars: 2mm.

### 2.6. *In Vivo* Assessment of Brain Foreign Body Response

Immunohistochemical analysis was performed on sections of neocortex collected ∼400-700 μm below the cortical surface of adult Sprague Dawley rats sacrificed 3 and 9-weeks post-implantation with MMBIs, PDMS implants, and silicon implants to assess the effects of implant stiffness on the FBR. The proteins Iba1, CD68, GFAP, and NeuN were selected for staining. Iba1 is distributed in the cytoplasm and processes of microglia and was used to assess microglia density and distribution. CD68, a transmembrane protein with increased expression in activated microglia, provided a distinction between activated and ramified microglia.^[35]^ GFAP, an intermediate filament protein upregulated in reactive astrocytes, was used to assess reactive astrocyte density and distribution.^[36]^ Finally, NeuN, a protein localized in the nuclei of neurons, was used to assess neuronal density.^[37]^ As with previous studies comparing the elicited FBR from implants made of stiffer materials such as tungsten or silicon to implants made of more compliant polymers such as Parylene-C or polyimide, there are differences in the chemical composition of each implant that cannot be excluded as a possible contributor to the change in the elicited FBR. Hence, in this study, the effects of the chemical differences were minimized between implants made from PDMS and Ecoflex which share many chemical attributes. The *in vivo* studies were evaluated at 3 and 9-week timepoints to compare the FBR elicited by the MMBIs, PDMS implants, and silicon implants shortly after the initiation of glial scarring and when the glial scar begins to mature.^[38]^

Representative confocal images of tissue sections with the implants removed are shown in **Figure 6a**. The same representative images with the individual stains can be seen in the supplementary Figures S7 and S8. The implant holes reflect the profile of the respective implants. Deviations from the expected shape and dimension could have resulted from processes such as tissue mounting, removal of the implants from tissue samples, or cryosectioning. Tissue sections with significant deviations (Figure S9, SI) were excluded from the study. We observed that Iba1, CD68, and GFAP immunohistochemical stains were localized around the tissue-implant interface and the intensity of the labeling decreased with increasing distance from the tissue-implant interface. Furthermore, Iba1, CD68, and GFAP intensity levels in tissue regions beyond 200 µm from the tissue-implant interface reflected the intensity levels observed in healthy tissue regions > 1 mm from the tissue-implant interface. Iba1, CD68, and GFAP intensity levels were quantified with respect to distance from the tissue-implant interface. Region of interests (ROI) were defined by drawing a rectangular outline with rounded corners around the implant hole. Subsequent outlines of the same shape were drawn in 10 µm increments until 400 µm away from the tissue-implant interface. An ROI bin consisted of the signal between two consecutive outlines (Figure S10, SI). Relative intensities presented in Figure 6b were obtained by normalizing the average intensity within a bin to the average intensity of the same tissue 700-900 µm from the tissue-implant interface. The relative intensity levels of Iba1, CD68, and GFAP within 50 µm from the tissue-implant interface were assessed for statistical significance using the non-parametric Kruskal-Wallis test (**Table 1**). 5 rats were used for each timepoint (n = 5). The specific number of tissue section replicates per rat for glial and neuronal stains are shown in Table S2. For statistical analysis, study conditions with multiple tissue section replicates had the results averaged and treated as a single measure.

**Table 1.**
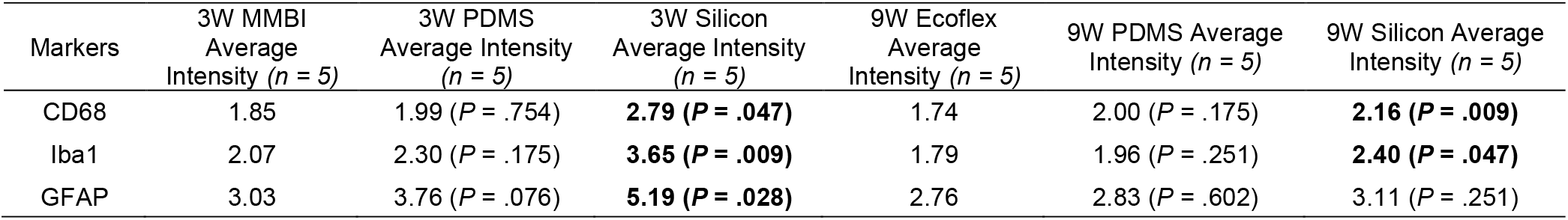
Comparison of CD68, Iba1, GFAP relative intensities 50 µm from tissue-implant interface for MMBIs, PDMS implants, and silicon implants 3 and 9-week post-implantation. P-values are calculated with respect to MMBIs and p-values < .05 are bolded.

**Figure 6.**
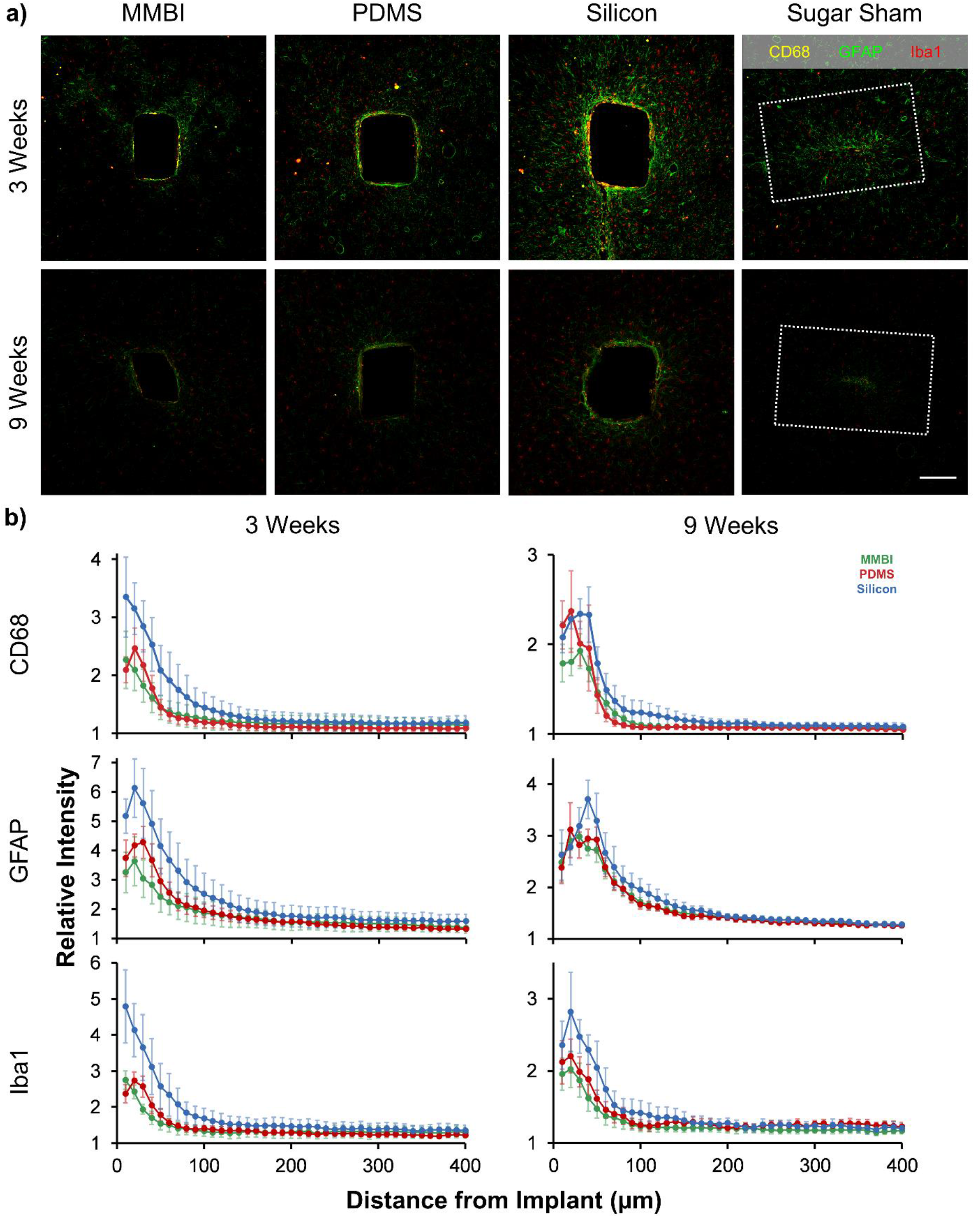
FBR elicited by MMBIs, PDMS implants, silicon implants, and sugar shams (shuttles with no encased implant). a) Representative confocal images of CD68 (yellow), GFAP (green), and Iba1 (red) elicited by the implants 3 and 9-week post-implantation. Dotted white lines in sugar sham images represent initial injury size. Images from each timepoint have identical image settings. b) Immunolabeling for CD-68, Iba1, and GFAP presented as normalized intensity curves with respect to distance from tissue-implant interface. Lower relative intensity signifies less brain FBR. Sugar shams were not analyzed due to insufficient number of implant tracks found. Error bars represent standard error. Scale bar: 200 µm.

The differences in relative intensity levels between MMBIs and PDMS implants did not meet the criteria of statistical significance (p-value < .05). Conversely, when comparing MMBIs to silicon implants, the differences in relative intensity levels met the criteria of statistical significance except for GFAP at 9-week post-implantation. However, in all instances, the average relative intensity levels within 50 µm from the tissue-implant interface were consistently lower for MMBIs compared to PDMS and silicon implants. We acknowledge that immunoreactivity detected in different tissue sections can introduce variability due to inherent variability in immunolabeling and the heterogeneity of cell populations. Addressing the impact MMBIs on specific neuronal and glial subpopulations in different brain regions will require additional studies.

All implants in this study were untethered, hence the observed FBR is not affected by external strain that arise in tethered implants and can be directly attributed to the interaction between the glial cells and the implant. Despite the absence of significant tissue strain arising from the displacement of brain tissue relative to the implant, the MMBIs elicited a lower FBR compared to both PDMS and silicon implants. It is also interesting to note that for sugar shams, implantation sites showed minimal inflammation at 3-week post-implantation and became almost indiscernible from healthy tissue at 9-week post-implantation. This observation is congruent with previous reports,^[39]^ supporting the conclusion that stab wounds are not the cause of chronic brain FBR.

The effect of implant stiffness on neuronal density was assessed by confocal microscopy of NeuN stained tissue sections and by calculating for neuronal density. The neuronal density of healthy tissues 400 µm from the tissue-implant interface were used as the baseline. Representative images of the neuronal density surrounding MMBIs, PDMS implants, and silicon implants are illustrated in **Figure 7a**. The neuronal density beyond 175 µm from the tissue-implant interface was like that of healthy tissue for all implants. Hence, we quantified neuronal density as a function of distance from the tissue-implant interface up to 200 µm away (Figure 7b). As with the glial labels, a rectangular outline with rounded corners was first drawn around the implant hole, but this time subsequent outlines were drawn in 25 µm increments up to 200 µm away. The signal between two consecutive outlines formed an ROI bin. The normalized neuronal density of each ROI bin was calculated, and as expected, the neuronal density increased with increasing distance from the tissue-implant interface. In all cases, neuronal loss was highest within 75 µm from the tissue-implant interface. Non-parametric Kruskal-Wallis tests were conducted for statistical significance. The difference in neuronal density from 25-50 µm (neuronal density from 0-25 µm was not analyzed because cell count was too low and variable) from the tissue-implant interface of MMBIs compared to PDMS and silicon implants met the criteria of statistical significance (p-value < .05) for both 3 and 9-week post-implantation. However, only the neuronal density surrounding MMBIs and PDMS implants at 3-week post-implantation met the criteria of statistical significance for 50-75 µm from the tissue-implant interface (Figure 7b). Longer duration *in vivo* studies will be needed to assess if the observed FBR benefits extend beyond the 9-week timepoint, and studies on neuronal health and activity in additional to neuronal density should be conducted in future work.

**Figure 7.**
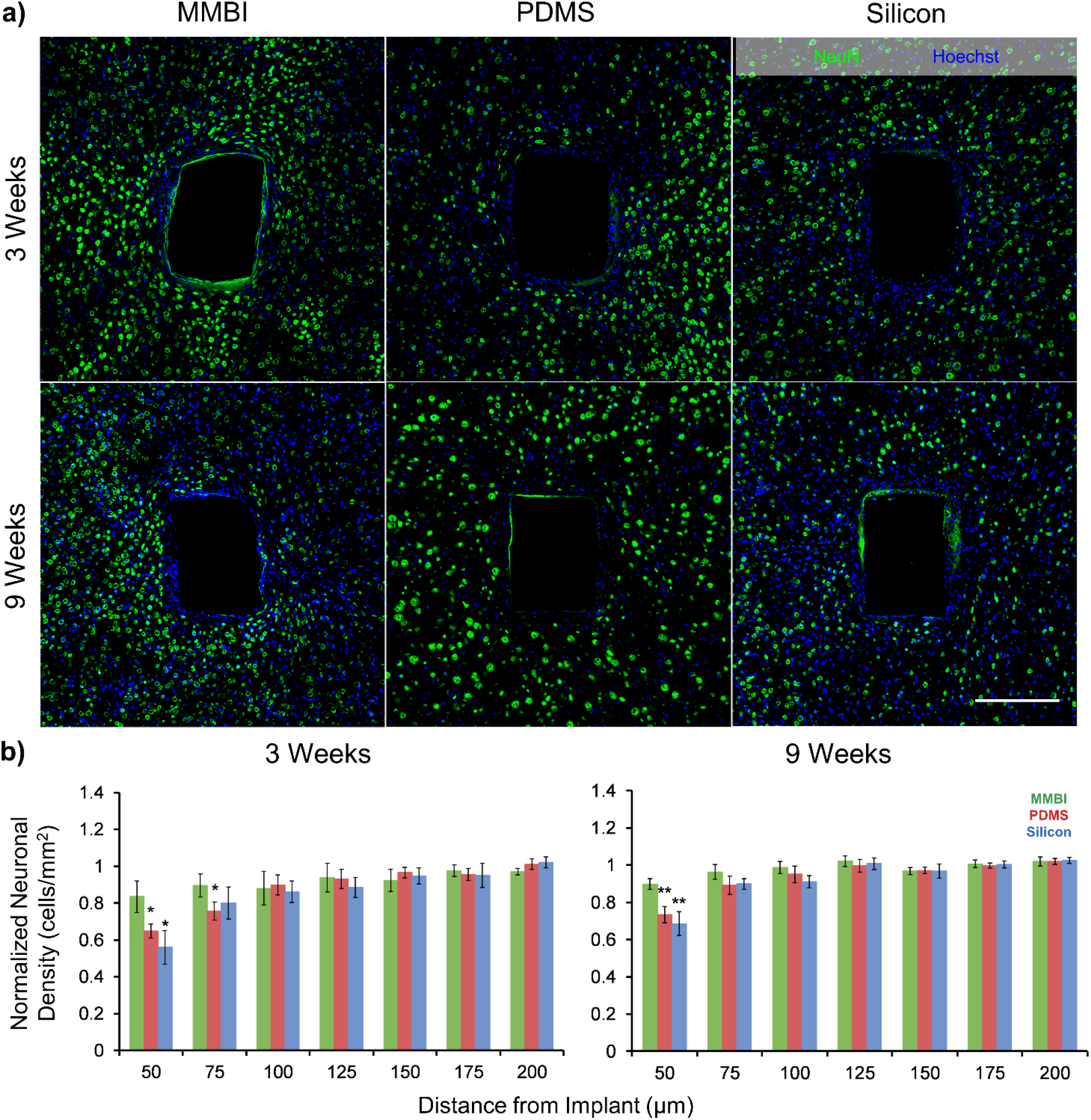
Neuronal density surrounding MMBIs, PDMS implants, and silicon implants. a) Representative confocal images of NeuN with Hoechst surrounding the tissue-implant interfaces 3 and 9-week post implantation. Sugar shams were not imaged and analyzed due difficulties in finding implants tracks and separating implant tracks from healthy tissue. b) Normalized neuronal density comparisons of PDMS and silicon implants with respect to MMBIs. * signifies significance at P < .05. ** signifies significance at P < .01. Actual p-values shown in Table S3 of the SI. N = 5 for all neuronal stains. Error bars represent standard error. Scale bar: 200 µm.

Electrical signals from neurons can be detected reliably up to 50-140 µm from the tissue-implant interface.^[40]^ Consequently, the presence of a substantial glial scar or a significant neuronal loss surrounding the tissue-implant interface could directly affect implant functionality and reliability. Based the observed FBR reduction and higher neuronal density in proximity to the MMBI tissue-implant interfaces, functional MMBIs may provide superior recording and better long-term reliability compared to conventionally stiff implants. However, future work is required to develop electrically functional MMBIs which is a prerequisite to benchmarking the electrical signal transduction of MMBIs to current, stiffer, electrically functional implants.

## 3. Conclusion

In summary, we introduced micrometer sized MMBIs (∼20 kPa) made by VAM that were encapsulated in dissolvable sugar shuttles and successfully and reliably delivered them into rat brains. The comparatively large sugar shuttles elicited minimal brain FBR. Our *in vivo* results comparing the FBR caused by MMBIs, PDMS implants, and silicon implants confirm and extend previous findings that show reduced FBR with increasing implant softness. MMBIs consistently elicited a reduced level of activated microglia, reactive astrocytes, and neuronal loss compared to PDMS and silicon implants at both 3 and 9-week post-implantation. VAM is versatile and could be adapted to develop functional MMBIs in the future by infilling conductive polymers or liquid metals into MMBIs with either embedded or open channels to make conductive traces. Furthermore, VAM coupled with sacrificial molds has the potential to create smaller MMBIs as the implant dimensions are determined by the master mold fabricated using photolithography. MMBIs could potentially improve long-term functionality and reliability of brain implants by minimizing strain and stress due to movements and swelling of the brain in both lateral and axial directions relative to the MMBI and help drive clinical adoption of brain implants.

## 4. Experimental Section

### Sacrificial Sugar Mold Fabrication

MD700 (Fluorolink MD700, Solvay) molds were replicated from SU-8 molds (Figure S1, SI) by pouring a mixture of MD700 and 2% (w/w) of a photo initiator (Darocur 1173, Sigma). The mixture was degassed on top of the SU-8 molds for 1 h and then UV cured (IntelliRay 600, UViTron International) for 2 min at 180 mW/cm^2^. The released MD700 molds were sonicated in isopropanol (IPA) for 15 min and then dried using a nitrogen gun. A 10:1 ratio of PDMS base and crosslinker (SYLGARD 184, DOW) was mixed and degassed using a planetary centrifugal mixer (AR-100, THINKY) at 2000 rpm for 90 s. The mixture was poured onto the MD700 molds and cured for 24 h at 60 °C (all PDMS were processed with these specifications unless otherwise stated). The released PDMS molds were placed feature-side into a silicone beaker containing corn syrup, water, and sucrose in a 1:2:3 ratio by volume and microwaved until the sugar mixture reached 180 °C. Afterwards, the container was quickly transferred to a vacuum oven set to 240 °C (Lindberg/Blue M, Thermo Scientific) and degassed at 25 inHg for 5 min. Once finished, the silicone container was left to cool at RT. PDMS molds were released from the solidified sugar resulting in the sugar molds. The sugar molds were stored in a vacuum desiccator until use.

### Vacuum-Assisted Molding of MMBIs

Sugar molds were sealed with an HRT (319Y-4LS, Nitto) exposing only the mold inlet and a PDMS O-ring was placed around the inlet. An Ecoflex mixture (Ecoflex 00-20, Smooth-On) of part A and part B in a 1:1 ratio (w/w) was mixed and degassed using a planetary centrifugal mixer at 2000 rpm for 90 s along with 10% of a silicone thinner (Silicone Thinner, Smooth-On) and .001% of a green silicone pigment (Silc Pig, Smooth-On) by weight. The mixture was quickly placed on ice afterwards and pipetted into the O-ring using a positive displacement pipette. The molds were then placed inside a vacuum desiccator and degassed for 20 min. After venting the desiccator, the Ecoflex prepolymer infilled the molds. The molds were placed in an oven at 60 °C for 1 h. The HRT was removed and the cured MMBIs were released by dissolving the molds in a water bath. MMBIs were then trimmed into 2.5 mm long pieces using a razor blade. For PDMS and silicon implant fabrication process see Figure S2 and S4 of the SI.

### Encasing Implants in Dissolvable Sugar Shuttles

Master molds were printed using a DLP printer (MiiCraft 100, MiiCraft) and a photocurable resin (BV-002A, MiiCraft). 50 μm layer thicknesses were used at an exposure of 0.8 s per layer. The 3D-printed molds were sonicated in IPA for 10 min and then post-cured in an UV flood curing system for 2 min at 180 mW/cm^2^. Afterwards, the molds were transferred to an oven and baked at 60 °C for 24 h to evaporate any uncured resin that inhibits PDMS curing. A PDMS mixture (10:1) was poured onto the 3D-printed molds and cured to create the PDMS molds. A layer of powdered sugar was deposited onto the features of the molds to prevent the MMBIs from sticking. A Kimwipe dipped in 100% ethanol was used to remove any excess sugar from the surface of the molds, and all implants were manually inserted using a tweezer. The molds were then sealed with a PDMS slab (20:1) that had a cut-out for the mold inlet and were dry autoclaved for 45 min at 121 °C. A sugar mixture (identical to the mixture used for sugar molds) was microwaved to ∼150 °C in a silicone beaker, and the PDMS molds were placed into the molten sugar with the inlet facing downwards. The beaker was quickly transferred to a vacuum oven set to 240 °C and degassed at 25 inHg for 20 min. Once the oven was vented, the sugar infilled. The molds were then quickly removed from beaker and left to cool at RT in a biosafety cabinet. The solidified sugar shuttles were released under sterile conditions using a tweezer and then stored in a vacuum desiccator until use.

### Surgical Implantation

Surgical procedures were done in accordance with the Canadian Council on Animal Care guidelines for the use of animals in research and were approved by the Montreal Neurological Institute Animal Care Committee. Stereotaxic surgery was performed on 10 female rats (Sprague Dawley Rat, Charles River) 10-12 weeks of age weighing 275-300 g to evaluate the *in vivo* brain FBR. For surgery specifics, see SI. Each rat had an MMBI, PDMS implant, silicon implant, and sugar sham implanted. Sugar shuttles with encased implants were mounted on an electrode holder (Model 1770, KOPF) and inserted until the encased implant could be seen entering the brain tissue completely. This was visually confirmed using an M50 microscope (M50, Leica) during surgery. Sugar shuttles were then left to dissolve. Sterile PBS was applied to the implantation site shuttles to facilitate dissolution of any remaining sugar and the incision was sutured after surgery. 5 rats were used per timepoint to evaluate the FBR. All rats at the 3-week timepoint developed superficial wound infections and were treated with subcutaneous injections (10 mg/kg body weight) of Baytril and wounds were cleaned daily for 5 days from the onset of infection. For MRI experiment protocol see SI.

### Histology

Rats sacrificed at the designated timepoints were anesthetized with intraperitoneal injection of Avertin (240 mg/kg body weight) and then transcardially perfused with ice cold PBS followed by 4% paraformaldehyde in PBS (w/v). Silicon implants were carefully removed from the brains before further processing. Explanted brains were cryoprotected in 30% sucrose prior to freezing, and 20 µm tissue sections were cut using a cryostat and mounted on microscope slides. For immunohistochemical analysis, two groups of co-staining were performed with the respective concentrations and concentrations listed in **Table 2**. Tissue sections ∼400-700 µm deep were used for analysis. Tissue sections were hydrated in 1X PBS and subsequently incubated in blocking solution (0.3% Triton X-100, 10% FBS, and 3% BSA in 1X PBS) for 2 h at RT. Afterwards, old blocking solution was removed and primary antibodies in new blocking solution were added to the tissue sections and incubated at 4 °C overnight. After primary antibody incubation, tissue sections were washed with 1X PBS three times for 10 min each. Secondary antibodies were diluted in new blocking solution and added to the tissue sections and incubated for 2 h at RT. After secondary incubation, tissue sections were washed with 1X PBS for 15 min followed by staining with Hoechst in 1X PBS for 15 min. Two additional 15 min washes in 1X PBS were performed before tissue sections were air dried and mounted (Fluoro-Gel, Electron Microscopy Sciences).

**Table 2.**
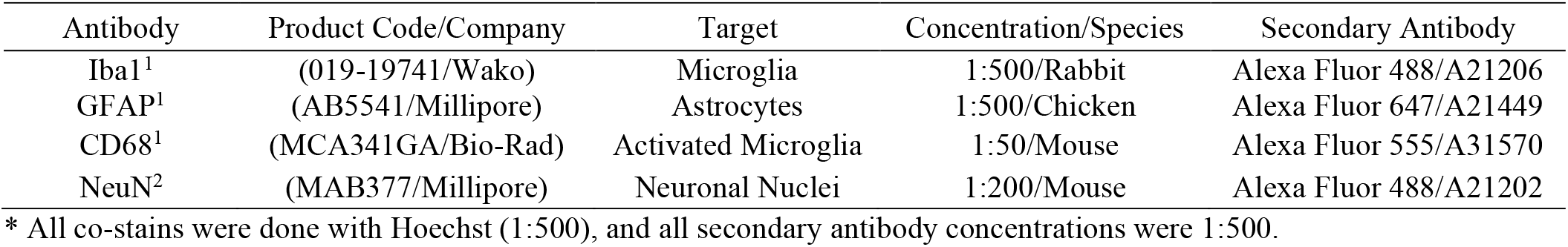
Table of antibodies used for immunohistochemistry brain FBR analysis.

### Tissue Analysis

Confocal fluorescent microscopy (Eclipse Ti2, Nikon) was performed on tissue sections to assess the brain FBR. For each antibody target, images of tissue sections from the same timepoint were acquired using the same laser power and detector settings to reduce variability. Laser power was adjusted such that there was less than 10% oversaturation in the pixels surrounding the tissue-implant interface. Multi-channel images were acquired simultaneously at 20X for glial stains and 40X for neuronal stains and subsequently stitched. Glial analysis was done using the MATLAB script I.N.T.E.N.S.I.T.Y v2.0 by Kozai *et al*.^[41]^ For each tissue section, tissue 700-900 µm from the tissue-implant interface was used as control. To calculate the background noise intensity for each image, the mean fluoresce intensity of the control area was first acquired and pixels one standard deviation above the mean fluoresce intensity value, which were considered signal, was removed from the calculation of background noise intensity threshold. The background noise intensity threshold was then set to one standard deviation below the mean fluoresce intensity value of the remaining pixels. Pixels below the background noise intensity threshold were considered holes, such as blood vessels, and were disregarded from calculations.

Implant holes that preserved the rectangular profiles of the implants were analyzed, and ROIs were first defined by drawing a rectangular outline with rounded corners around the implant hole. Subsequent outlines of the same shape were drawn in 10 µm increments until 400 µm away from the tissue-implant interface. An ROI bin consisted of the signal between two consecutive outlines (Figure S10, SI). The first outline was drawn slightly within the tissue-implant interface to capture all available signal. The intensity value of all pixels above the background noise threshold for each bin was averaged and normalized against the background generating a relative intensity profile as a function of distance from the tissue-implant interface. The normalized intensity values were calculated using **Equation 1** where *AvgI* is the averaged intensity value of all pixels above the background noise threshold in each bin, and where *AvgN* is the background noise threshold. For neuronal analysis, ROIs were defined the same way as the glial analysis, but outlines were drawn in 25 µm increments up to 200 µm away from the tissue-implant interface. The neuronal density was counted manually and was normalized to the neuronal density 400 µm from the tissue-implant interface. Results for the neuronal density and glial intensity levels from all tissue sections were averaged for each implant type and reported as the mean ± the standard error. Non-parametric Kruskal-Wallis tests were performed to determine statistical significance of the glial and neuronal results.

## Supporting Information

Supporting Information is available from the Wiley Online Library or from the author.

## Supporting information

MMBI Implantation in Rat

## Acknowledgments

The authors acknowledge funding from the National Science and Engineering Research Council of Canada (RGPIN-2016-06723, CHRP-493633-16), the Canadian Institute of Health Research (CHRP-357055), and the McGill University Healthy Brains for Healthy Lives program (1c-II-3). E.N.Z would like to thank McGill Nanotools and Polytechnique Montréal’s Laboratoire de Microfabrication for training in microfabrication as well as CMC Microsystems for access to SolidWorks and L-Edit used in this work. E.N.Z. would also like to thank Javier Alejandro Hernández-Castro and Auxtine Micalet for discussions and assistance in the VAM portion of the research. All experiments and data analysis were performed by E.N.Z. except for rat surgeries and cryosectioning performed by J.C., confocal microscopy performed by A.A., and MRI imaging performed by Marius Tuznik.

## Conflict of Interest

The authors declare no conflict of interest.

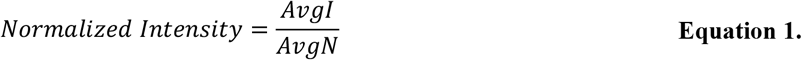

## Notes

### Competing Interest Statement

The authors have declared no competing interest.

