## Supplementary Material for "Mechanically Matched Silicone Brain Implants Reduce Brain Foreign Body Response"

**SU-8 Master Mold Fabrication**

**Figure S1** illustrates the master mold fabrication process. A 4 in prime grade silicon wafer (1115, UniversityWafer) was rinsed with acetone, isopropanol (IPA), and distilled water to remove organic residues. Afterwards, the wafer was dried using a nitrogen gun and placed on a hotplate (1000-1, Electronic Micro Systems) at 150 °C to dehydrate for 10 min. After dehydration, the wafer was centered onto a spin coater (WS-400, Laurell Technologies) and 5 ml of SU-8 (SU-8 2050, MicroChem) was poured onto the wafer in a spiral formation to avoid bubbles. The SU-8 was spun at 500 rpm for 5 s and then 1630 rpm for 30 s to achieve a thickness of 100 µm. Edge bead removal (EBR) was performed manually using a cotton swab dipped in SU-8 developer (MicroChem). The wafer was then soft baked at 65 °C for 5 min and 95 °C for 10 min and then cooled to room temperature (RT) on the hotplate. A second 100 µm layer of SU-8 was coated with the same spinning parameters but with longer soft bake times to account for the insulation of the first layer (65°C for 10 min and 95°C for 20 min). A 20k DPI transparency photomask (CAD/Art Services Inc.), shown in Figure S1b, was placed in proximity contact with the wafer at a separation distance of 100 µm and the wafer was exposed using a mask aligner (EVG620, EV Group) at 400 $mJ/cm^2$ to crosslink. After exposure, the wafer was left to rest for 10 min before post exposure baking at 65 °C for 10 min and 95 °C for 26 min and then cooled to RT on the hotplate. The wafer was then placed into SU-8 developer to dissolve the unpolymerized SU-8 for 7 min or until no more residual unexposed resist can be seen. The fabricated SU-8 master mold was rinsed with fresh SU-8 developer followed by IPA and then dried with a nitrogen gun.

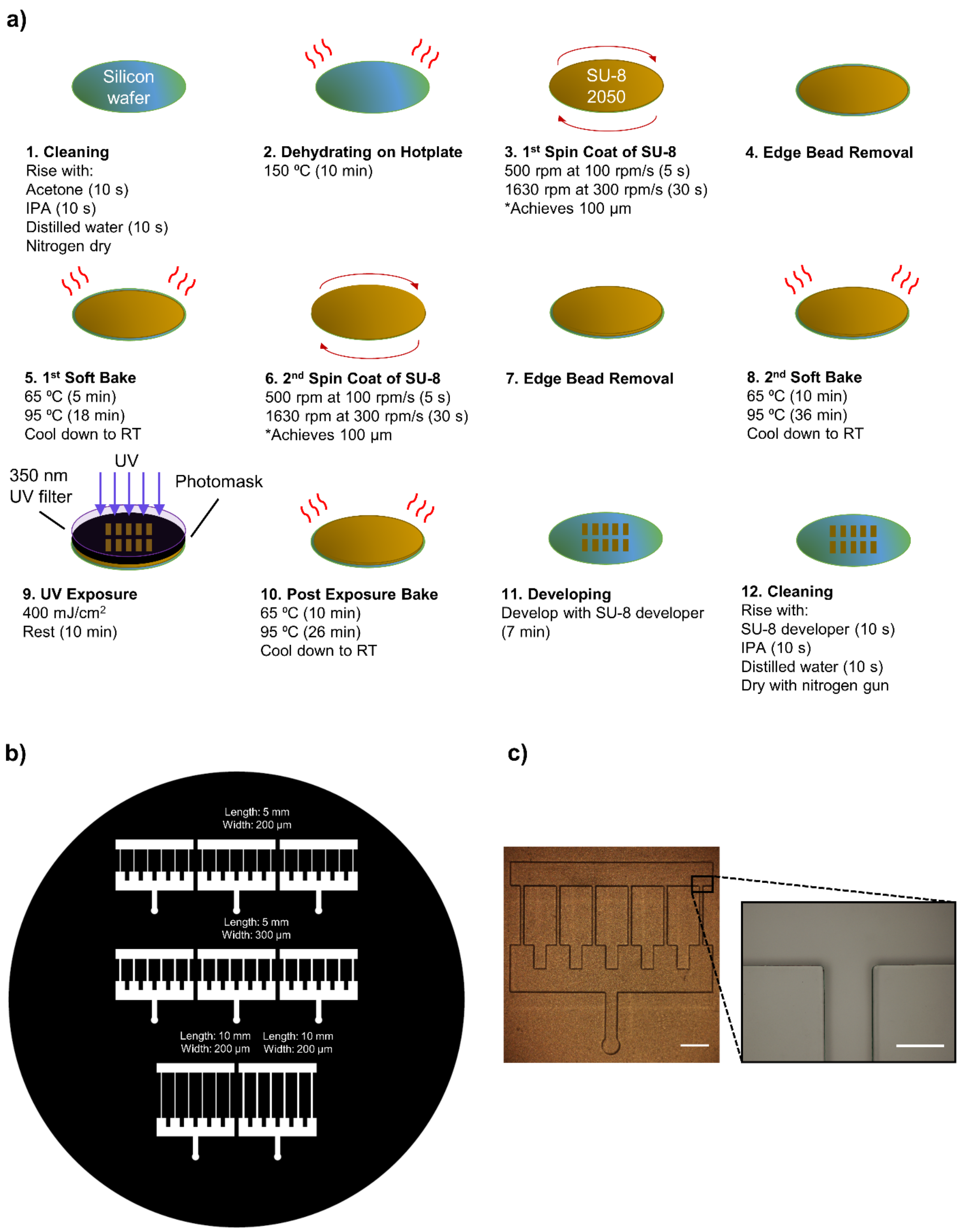


**Figure S1.** a) Fabrication process of SU-8 master mold. b) Photomask design for SU-8 master mold with feature dimensions. Four implant variants were made. c) Microscope image of negative MD700 mold. Scale bars: 2 mm, 300 µm (inset).

### PDMS Implant Fabrication

Negative MD700 molds were used for VAM of PDMS implants. See **Figure S2** for fabrication process. A different HRT from the Ecoflex VAM was used (REVALPHA 3198LS, Nitto). This tape adhered stronger to the PDMS implants and consequently released the PDMS implants from the MD700 molds more reliably. PDMS base and crosslinker in a 10:1 ratio was mixed using a planetary centrifugal mixer at 2000 rpm for 90 s and degassed before being pipetted using a viscous pipette into the PDMS ring on the MD700 molds. The molds were then degassed for 20 min and then vented to allow the PDMS prepolymer to infill the molds. After infilling, the PDMS was cured at 60 °C for 24 hr. Once cured, removing the HRT from the MD700 molds also released the PDMS implants. PDMS implants were released from the HRT by heating the HRT to 90 °C on a hotplate.

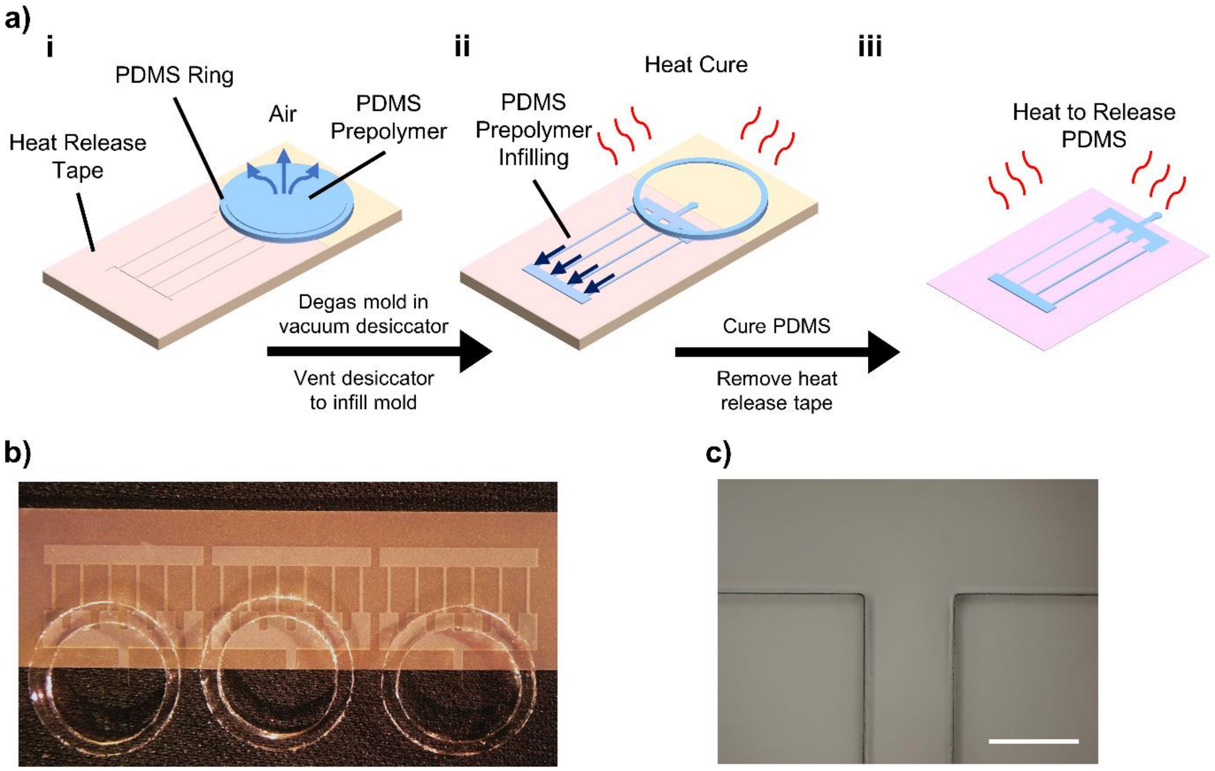


**Figure S2.** a) Fabrication process of PDMS implants. b) MD700 mold setup for VAM. c) Microscope image of PDMS implant. Scale bar: 300 µm.

### Silicon Implant Fabrication

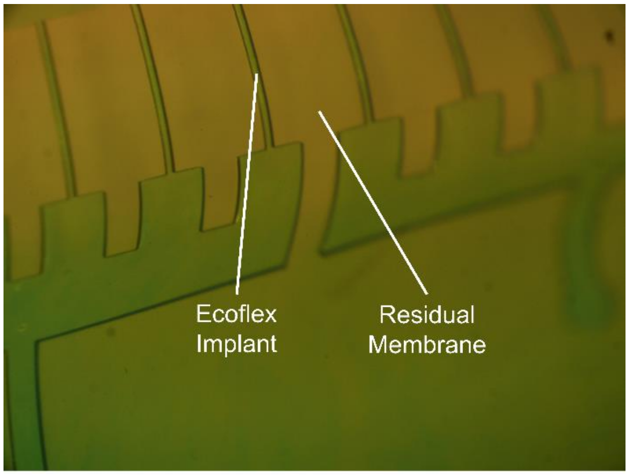


**Figure S3.** Ecoflex implants with residual membrane connecting independent features.

Bosch etching was used to fabricate the silicon implants. A protective mask (**Figure S4**) was created using a 2 in prime grade silicon wafer (200 µm thick) cleaned with acetone and IPA followed by distilled water. The wafer was dried using a nitrogen gun and baked on a hotplate at 150 °C for 10 min. Afterwards the wafer was spin-coated with SU-8 negative photoresist (SU-8 2015, MicroChem) at 500 rpm for 5 s and then 2000 rpm for 30 s using a spin coater to achieve a thickness of 20 µm. EBR was performed using a Q-tip dipped in SU-8 developer. Afterwards, the wafer was soft baked at 65 °C for 2 min and 95 °C for 4 min and then cooled to RT on the hotplate. A 10k DPI transparency photomask was placed in proximity contact with the photoresist and was exposed using a mask aligner at 200 $mJ/cm^2$ to crosslink. Afterwards, the wafer was left to rest for 10 min at RT before a post bake at 65 °C for 2 min and 95 °C for 5 min and then cooled to RT on the hotplate. The wafer was then placed in SU-8 developer to dissolve the unpolymerized SU-8 for 2 min. The wafer was rinsed with fresh SU-8 developer followed by IPA and distilled water and dried with a nitrogen gun.

After the protective mask was created, the wafer was mounted onto a 6 in silicon carrier wafer with a mounting adhesive (Crystal Bond 555 HMP, Agar Scientific). Anisotropic Bosch etching was used with an 8:20 passivation to etching time. The polymer deposition step was as followed: 8 seconds, 65 sccm $C_4F_8$, 1 sccm $SF_6$, ICP power 450 W, platen power (CCP) 10 W. The etching step was as followed: 20 seconds, 65 sccm $SF_6$, 15 sccm $C_4F_8$, ICP power 450 W, platen power (CCP) 25 W. An etch rate of 700 nm/step (~1.4 µm/min) was achieved. After successful etching, the silicon carrier wafer was heated to 70 °C causing the mounting adhesive to flow and to release the etched silicon implants. Lastly, the silicon implants were sonicated in IPA for 20 min to remove any adhesive residue.

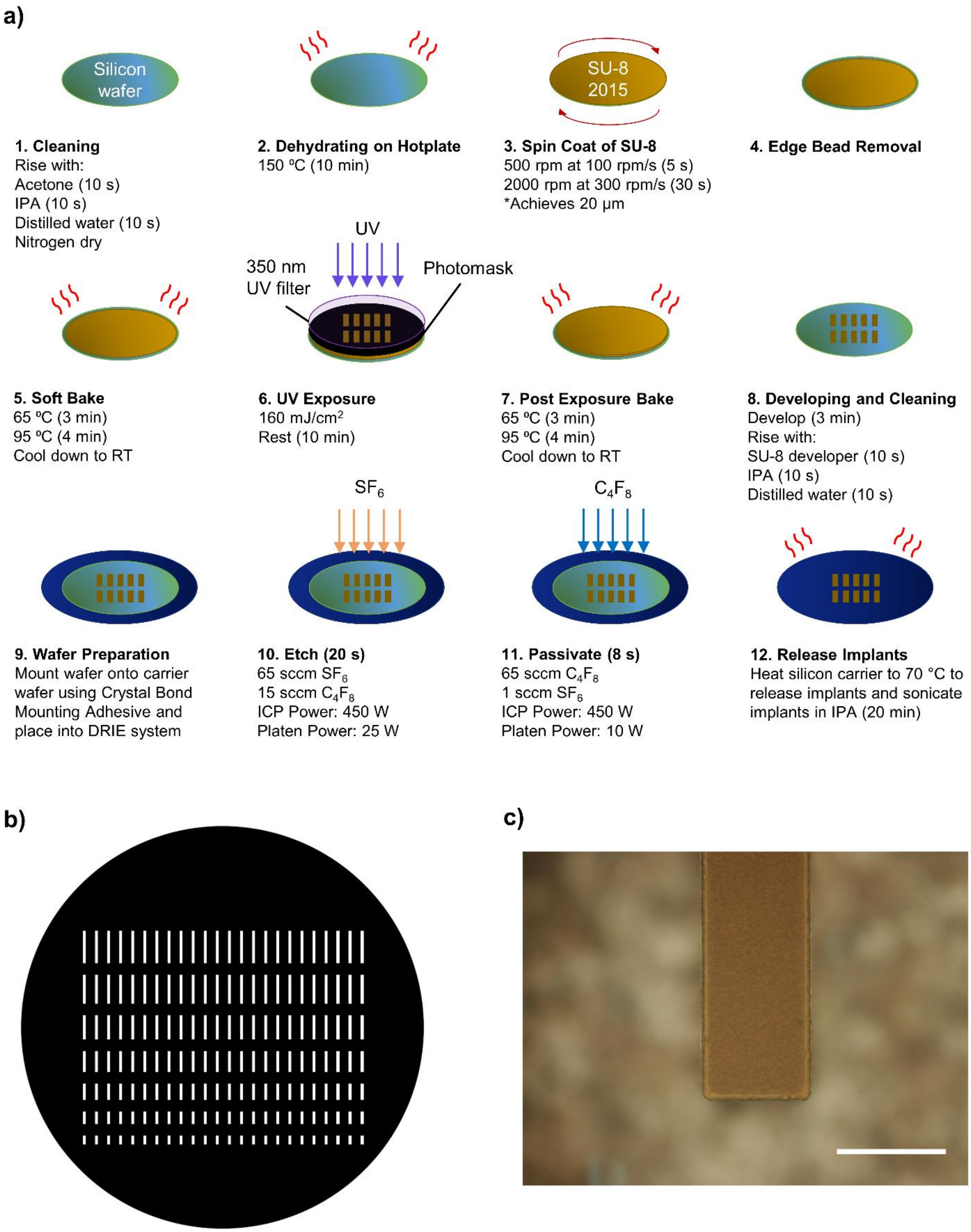


**Figure S4.** a) Fabrication process of silicon implants. b) Photomask design for SU-8 mask for silicon implant fabrication. All features were 300 µm wide ranging from 1 to 4 mm in length in .5 mm increments. c) Microscope image of silicon implant. Scale bar: 300 µm.

**Characterization of Ecoflex and PDMS Young's Modulus *(E)***

Tensile testing (5965 Series Universal Testing System, Instron) was performed on Ecoflex and PDMS test specimens to obtain the *E*. Ecoflex prepolymer consisted of part A and part B in a 1:1 ratio with an added silicone thinner that was 10% of the total weight of the Ecoflex prepolymer. The prepolymer was mixed using a planetary centrifugal mixer at 2000 rpm for 90 s for both mixing and degassing. PDMS prepolymer consisted of the base and crosslinker in a 10:1 ratio was mixed the same way as the Ecoflex prepolymer. The Ecoflex or PDMS prepolymer was poured onto an aluminum mold fabricated according to the ASTM D412 Type C specifications (**Figure S5**) to make the test specimens. Bubbles formed from pouring were removed with a pipette and the

a) b)

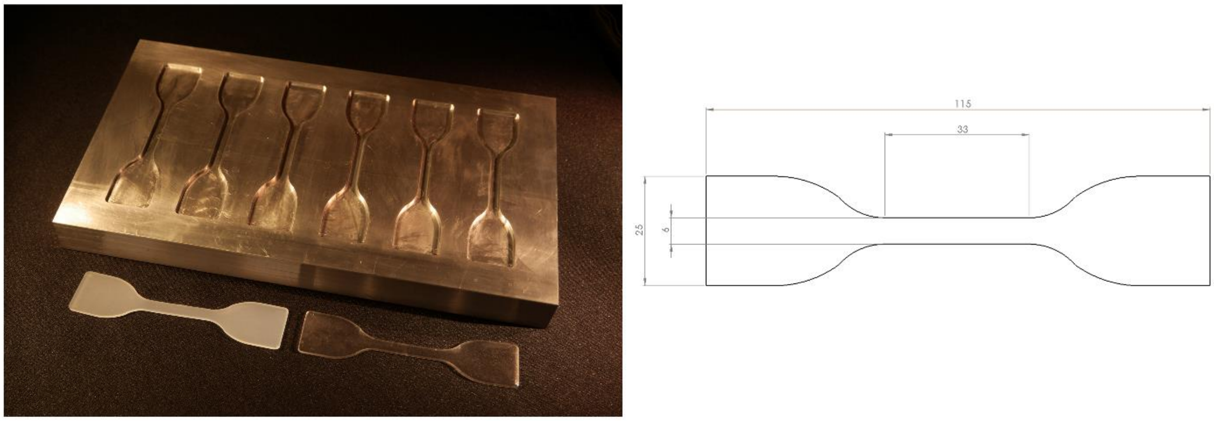


**Figure S5.** a) Aluminum mold with ASTM D412 Type C specifications used to make Ecoflex (bottom left) and PDMS (bottom right) test specimen. b) Dimensions of test specimen.

Ecoflex prepolymer was cured at 60 °C for 1 hr while the PDMS prepolymer was cured at 60 °C for 24 hr. An additional experimental condition in which the cured Ecoflex and PDMS specimens underwent a dry autoclave cycle for 45 min at 121 °C and a subsequent heat treatment in a vacuum oven for 20 min at 240 °C was conducted to investigate the effects of the sterilization and sugar encasing processes on the *E* of the Ecoflex and PDMS implants. The thickness of each test specimen was measured using a micrometer (P/N TL268, Proster) before being mounted onto the tensile machine. A 10 N load cell was used for Ecoflex and a 1 kN load cell was used for PDMS.

The crosshead velocity was set at 3 mm/s. Test specimens were extended until failure or until the tensile machine reached its extension limit. In total six test specimens were used to acquire the *E* of each experimental condition. All test data was obtained using Bluehill Universal. See **Figure S6** for the resultant stress-strain curves.

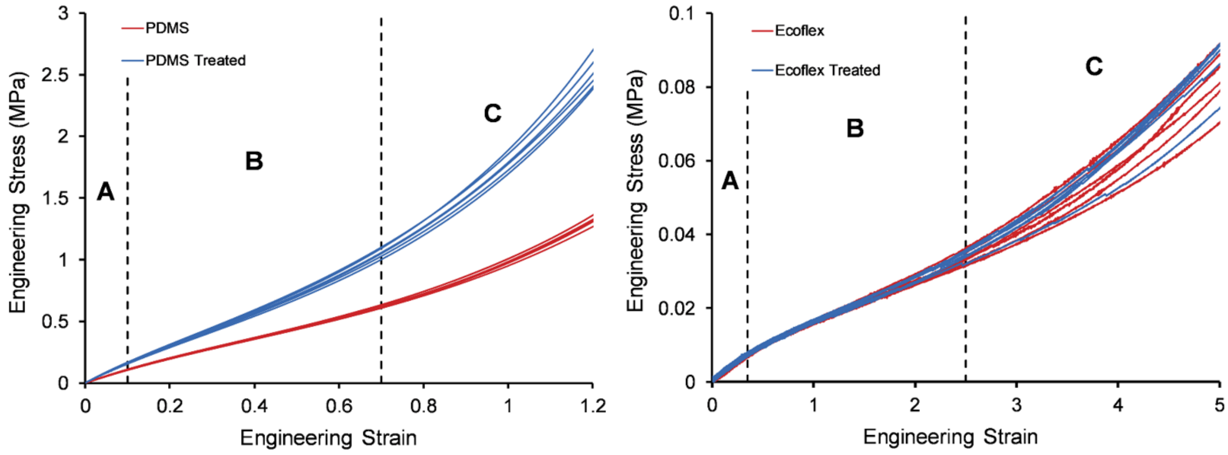


**Figure S6.** PDMS and Ecoflex stress-strain curve with (blue, n = 6) and without (red, n = 6) additional heat treatment show significant changes in PDMS (p-value < .05) but not in Ecoflex stiffness (p-value > .05). A, B, and C identifies regions of the stress-strain curves that show a distinct change in material stiffness with respect to strain. Linear elastic segments in region A were used to calculate the *E* of Ecoflex and PDMS with and without additional heat treatment.

Ecoflex and PDMS both exhibit typical stress-strain curves of hyper-elastic materials with three distinct regions depicted as A, B, and C where at strain rates within region A, the silicone elastomers are relatively stiff compared to strain rates in region B, followed by an increase in stiffness at strain rates in region C. Region A, the initial linear elastic segment of the stress-strain curve at low strains where Hooke's law is valid[1] was used to calculate the *E* of Ecoflex using **Equation 1** where σ is the applied stress and ε is the resultant strain.

**Equation (1).**

$$E = \frac{\sigma}{\varepsilon}$$

Since the implants are untethered to the skull after insertion into the brain, the only source of strain that is experienced by the implants is from brain tissue micromotions that results in tissue displacements of 2-30 μm in rats.[2] Given this relatively small displacement compared to the dimensions of our implants, the *E* obtained from region A serves as appropriate characterizations of the stiffness of our implants.

**Table S1.** Fracture strains of PDMS and Ecoflex. Ecoflex fracture strain limit was beyond 950% but was limited by the range of the tensile machine.

| Material | Fracture Strain | Fracture Strain with Additional Heat Treatment |
| --- | --- | --- |
| Ecoflex 00-20 | Tensile Limit (950%) | Tensile Limit (950%) |
| PDMS | 233% | 183% |

### Surgical Procedure for Rats

Surgical procedures were carried out in accordance with the Canadian Council on Animal Care guidelines for the use of animals in research and were approved by the Montreal Neurological Institute Animal Care Committee. Animals were anesthetized with 3% isoflurane in 0.8 L/min oxygen prior to surgery and maintained during the procedure at 2-2.5% isoflurane via a facemask. Ophthalmic ointment was applied to both eyes to prevent dryness and damage. Afterwards, 10 mg/kg body weight of carprofen (RIMADYL) and 0.5 mL/10 g body weight of 0.9% sterile saline was administered subcutaneously. Pedal reflex tests were performed before the beginning of each surgery and anesthesia level was monitored closely during the surgery by observing the breathing rate of the rats. Rats were placed on a stereotaxic (Model 940, KOPF) and four 1.5 mm holes were drilled using a handheld drill (Micromotor 1070, Foredom) into the skull at the coordinates where the implants were to be delivered (Figure S11, SI). Holes were drilled until a thin layer of skull was left and the remaining skull was removed manually using a 25G syringe with the tip bent into a hook shape to avoid damaging brain tissue. Coordinates were chosen to avoid major ventricular systems and such that there was no overlap in the evoked brain FBR.

**Magnetic Resonance Imaging of MMBI Delivery**

An adult rat was immediately sacrificed after the implantation of the MMBI and was anesthetized with intraperitoneal injection of Avertin (240 mg/kg body weight) and then transcardially perfused with ice cold PBS followed by 4% paraformaldehyde in PBS (w/v). The fixed brain was removed from the skull and placed in a large syringe filled with MR-inert fluid (Fluorinert, 3M). The syringe was then placed on the sample table at a distance coinciding with the center of the transceiver coil of a Bruker Pharmascan 7T MRI scanner. The scan was conducted using Paravision 5.1 with the following scanning parameters: 3D FISP, TR = 12 ms, TE = 6 ms, 4110.02 ms (scan repetition time), field of view = 3 cm x 1.8 cm x 1.41 cm, matrix size = 400 x 240 x 188, 75 µm isotropic voxels.

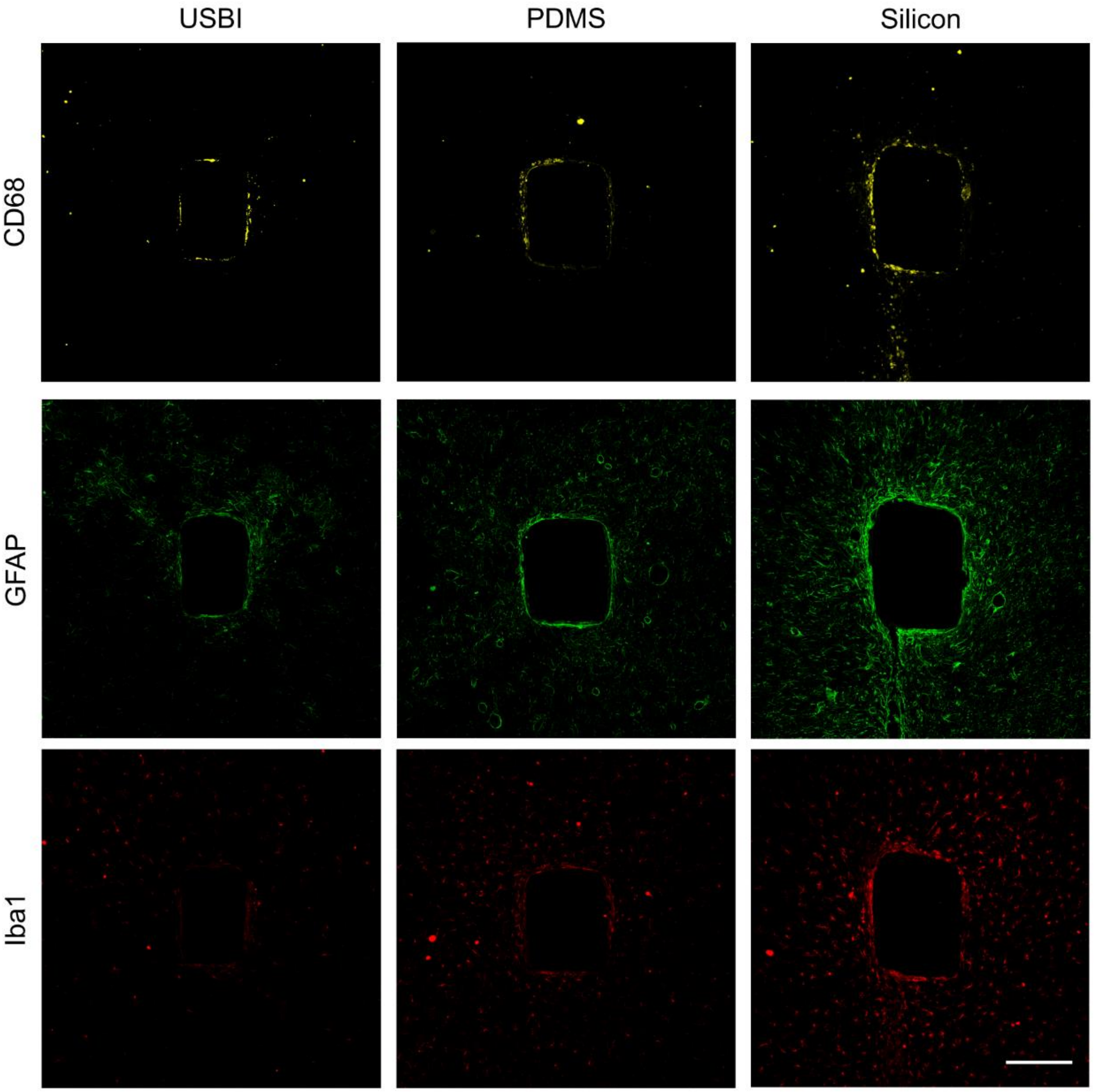


**Figure S7.** Representative confocal images of CD68 (yellow), GFAP (green), and Iba1 (red) elicited by the implants 3-week post-implantation. Scale bar: 200 μm.

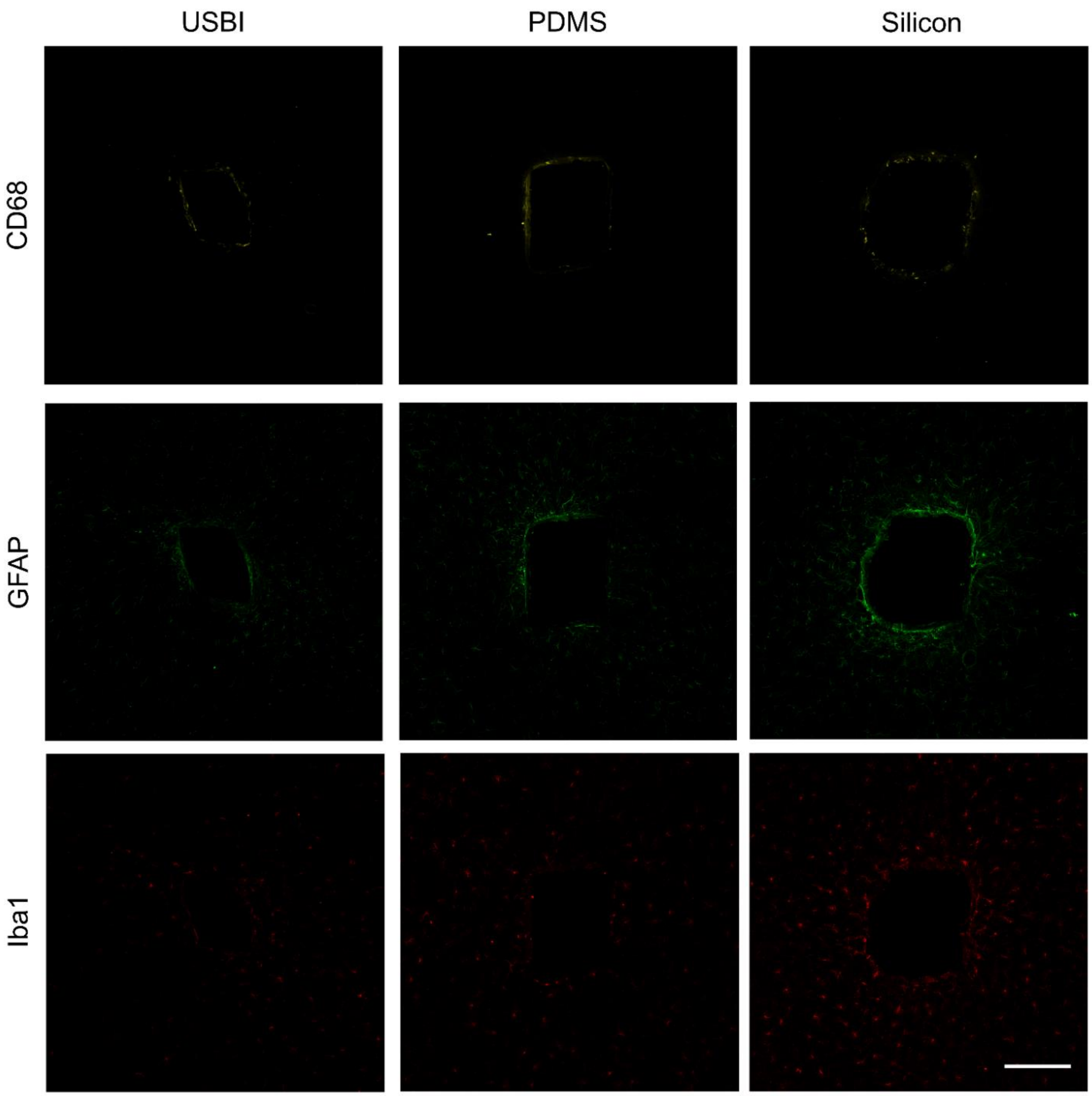


**Figure S8.** Representative confocal images of CD68 (yellow), GFAP (green), and Iba1 (red) elicited by the implants 9-week post-implantation. Scale bar: 200 µm.

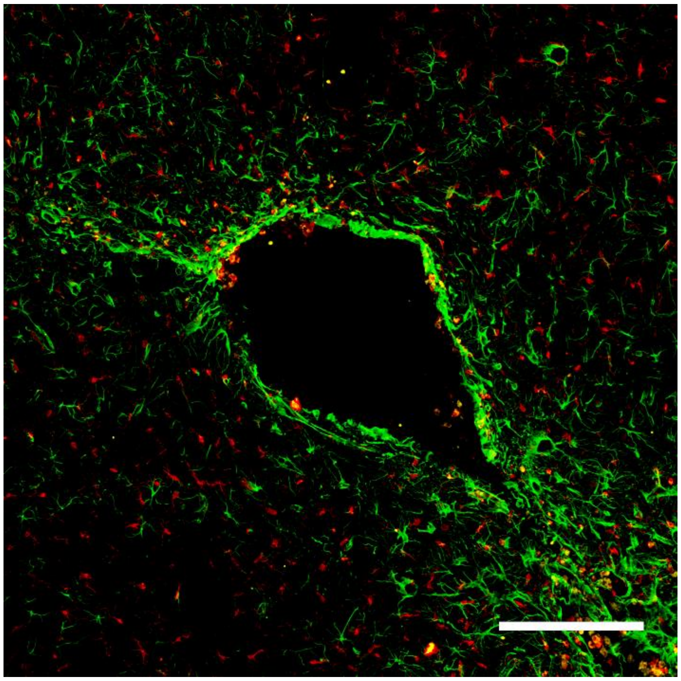

**Figure S9.** Example of implant hole irregularity. Samples such as these were not used for data analysis. Scale bar: 200 μm.

**Table S2.** Comparison of normalized average neuronal density 50 and 75 μm away from Ecoflex, PDMS, and silicon implants 3 and 9-weeks post-implantation. P-values given are with respect to a comparison with Ecoflex implants.

| Markers | 3W Ecoflex Average NeuN Density | 3W PDMS Average NeuN Density | 3W Silicon Average NeuN Density | 9W Ecoflex Average NeuN Density | 9W PDMS Average NeuN Density | 9W Silicon Average NeuN Density |
|---|---|---|---|---|---|---|
| NeuN at 50 μm | .83 | .65 (P = .046) | .56 (P = .012) | .90 | .73 (P = .002) | .69 (P = .002) |
| NeuN at 75 μm | .89 | .75 (P = .027) | .80 (Non-significant) | .96 | .89 (Non-significant) | .90 (Non-significant) |

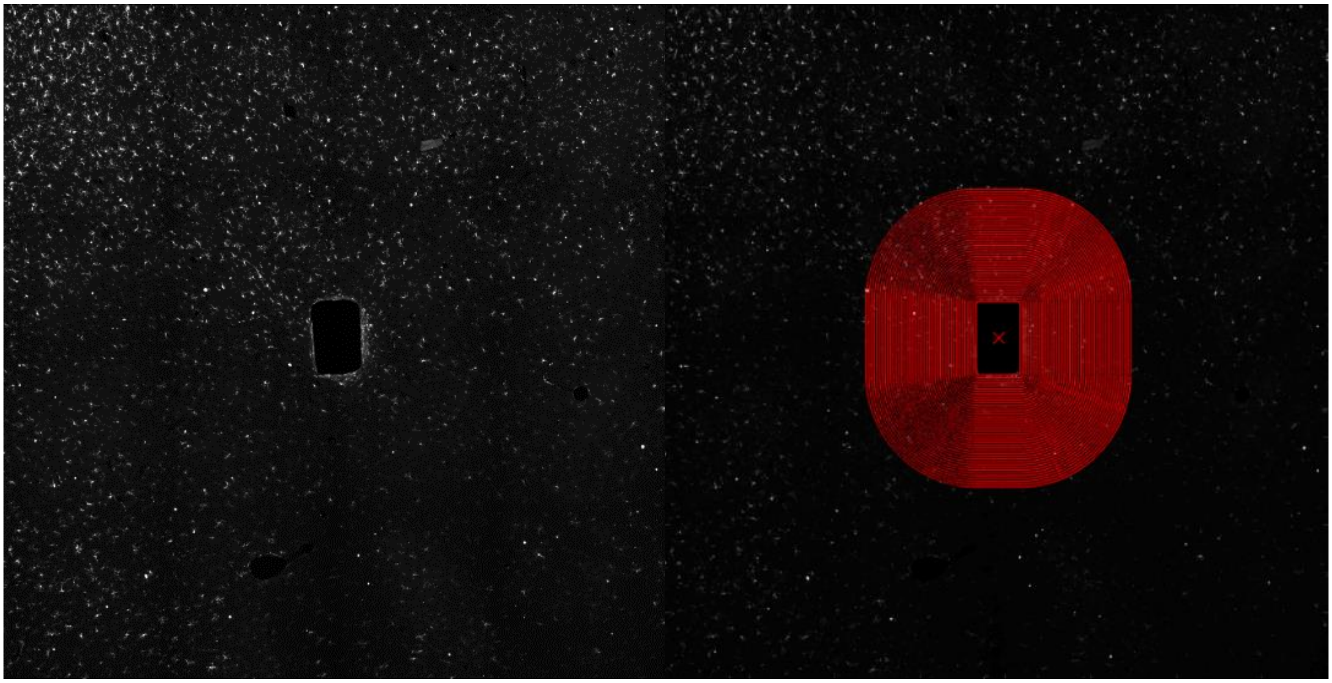

**Figure S10.** Rectangular regions of interest with 10 μm increments up to 400 μm away from the tissue-implant interface. Image shows Iba1 stain.

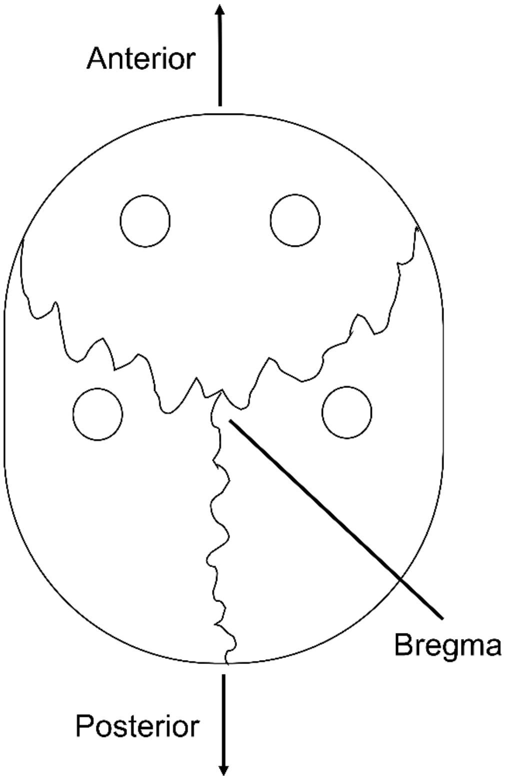


**Figure S11.** Coordinates of implantation sites relative to bregma. 2.5 mm anterior to bregma, 3 mm left and right. -.5 mm anterior to bregma 4 mm left and right.
